# Comparison of null models for combination drug therapy reveals Hand model as biochemically most plausible

**DOI:** 10.1101/409946

**Authors:** Mark Sinzger, Jakob Vanhoefer, Carolin Loos, Jan Hasenauer

## Abstract

Null models for the effect of combination therapies are widely used to evaluate synergy and antagonism of drugs. Due to the relevance of null models, their suitability is continuously discussed. Here, we contribute to the discussion by investigating the properties of five null models. Our study includes the model proposed by David J. Hand, which we refer to as *Hand model*. The Hand model has been introduced almost 20 years ago but hardly was used and studied. We show that the Hand model generalizes the principle of dose equivalence compared to the Loewe model and resolves the ambiguity of the Tallarida model. This provides a solution to the persisting conflict about the compatibility of two essential model properties: the sham combination principle and the principle of dose equivalence. By embedding several null models into a common framework, we shed light in their biochemical validity and provide indications that the Hand model is biochemically most plausible. We illustrate the practical implications and differences between null models by examining differences of null models on published data.

## 1. Introduction

Combination drug therapy is an advancing field of research in oncology, anesthesiology and immunology (Mao *et al.*, 2011; Dahl *et al.*, 2014; Bansal *et al.*, 2014; Hu *et al.*, 2016; Ahn & Root, 2017; Eser & Jänne, 2018). Treatments with multiple drugs are studied to distribute side effects and minimize toxicity while attaining the full efficacy (Maguire & France, 2018). The discovery of synergistic combinations can enhance the development of multi-drug therapies and was ranked second place in the principles of combination therapy after decreased “toxicity without decreased efficacy” (Levin & Harris, 1975). The term ‘combination’ refers to the choice of pairing agents *A* and *B*, to their specific concentration ratio as well as to their explicit doses. In the following we will use the terms drug and agent interchangeable.

Synergy is detected and quantified via the comparison of an experimentally obtained effect and the mathematical reference effect of a null model. If the measured effect to a combination therapy exceeds the reference effect based on the measured effects of the individual drugs, the dose pair is considered synergistic, otherwise antagonistic. In order to quantitatively assess the level of synergy, concepts such as combination indices (Chou & Talalay, 1984) or tools of statistical analysis (Tallarida *et al.*, 1989; Hennessey *et al.*, 2010) were introduced. A careful choice of the null model used to study synergy is important to not over-interpret results of drug combination studies (see, e.g., (Palmer & Sorger, 2017) for a study where drug independence was able to explain data rather than synergy or additivity). However, this choice requires a detailed understanding of the null models and the differences between them.

Applied in combination, the single effects of an *A*-dose and a *B*-dose are expected to add up in a way the null model specifies. Although there exist cases in which the simple algebraic sum of the effects may serve as a null model (Slinker, 1998), in general this sum will exceed the maximal possible effect, e.g., when the effect is measured in percentage of bound agents. A variety of null models has been introduced in the past and each model tries to provide a plausible concept of additivity that can be biochemically justified. Conceptually, effect-based strategies and dose-effect based strategies are distinguished. In the established Bliss model (Bliss, 1939), normalized effects are interpreted as probabilities of stochastically independent events, e.g., the event of an analgesic binding to a receptor. This effect-based strategy permits an additivity concept according to the addition rule of probabilities. Two agents which act on different components in a biochemical cascade can be modeled in this mutually non-exclusive way (e.g., the HIV-1 entry inhibitors CoRA and FI (Ahn & Root, 2017)). However, the implicit assumption of stochastic independence does not hold in case of drugs that act through a similar mechanism. For example, cetuximab and afatinib both inhibit EGFR in the pathway of non-small cell lung cancer proliferation (Moran, 2011; Janjigian *et al.*, 2014). In particular, a sham combination, i.e., a combination of an agent with itself, cannot be described by the Bliss model (Greco *et al.*, 1995).

Other null models incorporate knowledge on the functional relationship between dose concentration and effect for each single agent over a large range of doses. Such dose-effect based strategies shift the task of designing a null model towards specifying a plausible theory of addition of functions instead of a few numeric values (Geary, 2013). This functional relationship between dose concentration and effect for the mono-therapies are frequently modeled as Hill curves, which can be fitted to experimental data by standard software (Motulsky & Christopoulos, 2004). Here, we assume, that the dose-effect curves are increasing and have their maximal numerical value at the saturation level. However, minor changes allow an analogous treatment of decreasing dose-effect curves.

The classical dose-effect based model for mutually exclusive agents was first introduced by Loewe & Muischnek (1926); Loewe (1953) and popularized by Berenbaum (1977); Chou & Talalay (1984). It states that doses of drug *A* and drug *B* may be linearly converted into each other inducing no modification in effect, where the conversion rate generally depends on the current effect level. For example, if both drugs attain the same maximal effect, then one third of the half-max concentration of *A*, i.e., the concentration at which the half-max effect is reached, combined with two thirds of the half-max concentration of *B* is predicted to yield an observed effect of 50%. The set of all dose combinations in the ([*a*], [*b*])-plane, that are predicted to induce the same effect level is called an isobole, where [*a*] and [*b*] are the doses of *A* and *B*, respectively. Being an indicator of zero interaction, the isobole separates the plane into a region of synergy of points located below the isobole and a region of antagonism located above it. For the Loewe model isoboles are straight lines, whose slopes are the conversion rates as depicted by the isobologram in Figure 1A.

**Figure 1:**
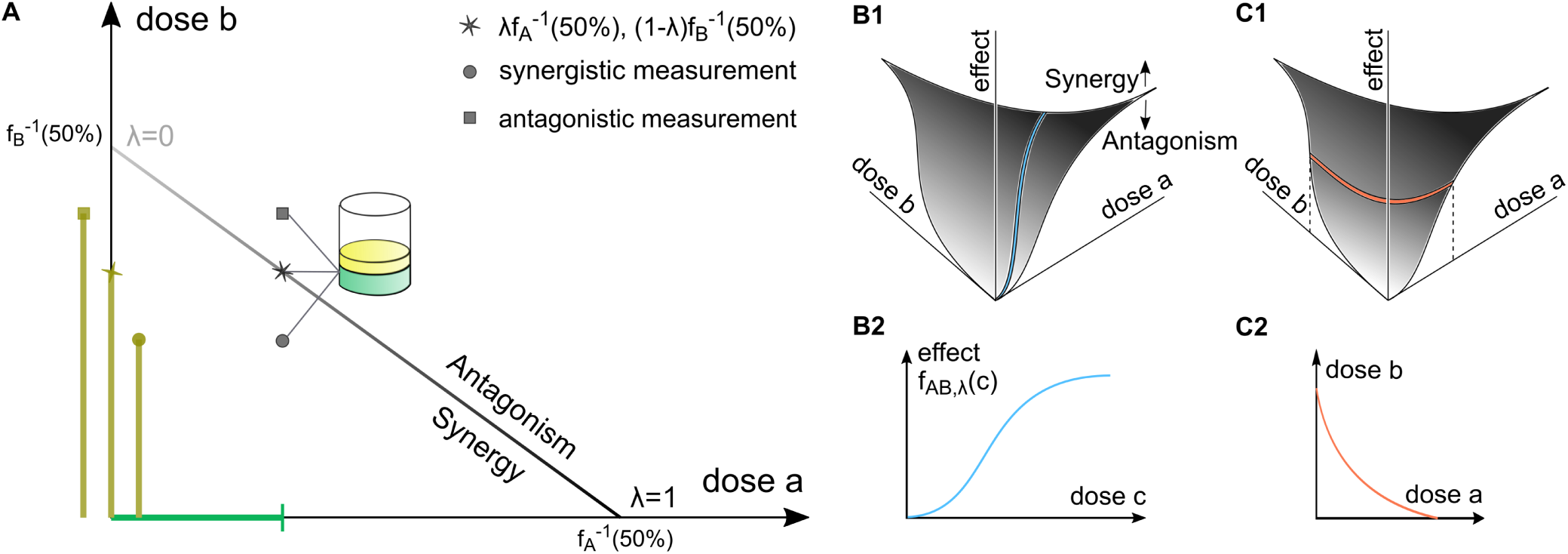
Loewe isobologram and the general effect surface. **(A)** Straight isobole at effect level 50% in the Loewe model. If the dose pairs located at star, circle, box induced 50%, the Loewe model would rate them non-interacting, synergistic and antagonistic, respectively. **(B)**-**(C)** visualize a general effect surface with informative cuts. **(B1)** Measured effects above and below the predicted effect *E* indicate synergy and antagonism, respectively. Vertical cuts through the effect surface along rays correspond to **(B2)** dose-effect curves of combined agents. **(C1)** Horizontal cuts through the effect surface provide **(C2)** isoboles.

The Loewe model is broadly accepted and used in cases of constant potency ratio. The potency ratio is the ratio of the effects of two drug and two dose-effect curves are said to have a constant potency ratio if they are identical up to rescaling the dose axis. This means that they are parallel in the representation with a logarithmic dose scale. We will refer to this central but rare case as the ‘constant potency ratio case’ or the ‘case of parallel dose-effect curves’. If the dose-effect-relations are given by Hill curves, two curves exhibit a constant potency ratio, exactly if they coincide in their maximal effects and Hill coefficients. In the prominent cases of a full and a partial agent or different cooperativities, i.e., maximal effect or the Hill coefficient differ, the potency ratio varies across effect levels.

The Loewe model is accepted for parallel dose-effect curves because this case implies that the conversion rate does not depend on the current effect level, i.e., the isoboles for all effect levels in the Loewe model are parallel. Two principles were extracted from the constant potency ratio case that seem to permit a transfer to the varying potency ratio case: The principle of dose equivalence (Grabovsky & Tallarida, 2004) and the principle of sham combinations (Berenbaum, 1989). The former states that each *A*-dose can be assigned a *B*-equivalent, which induces the same effect. The latter states that, a drug is not expected to synergize in a combination with itself. The sham combination principle is rather specific to mutually exclusive agents and violated by several null models, including the Highest Single Agent (HSA) model, the Bliss model and the Fisher’s dosage orthogonality (Russ & Kishony, 2017).

The attempt to base a null model on coupling both principles for curves of varying potency ratio (Grabovsky & Tallarida, 2004; Tallarida, 2006, 2011) fails to identify unique isoboles, as objected by Lorenzo & Sánchez-Marin (2006). This failure has incorrectly been attributed to Loewe as the caption “indeterminate Loewe additivity solution”, which Geary uses in (Geary, 2013), indicates. In fact, Loewe cannot be indeterminate, because the model, by definition, postulates a unique linear isobole. To the contrary, the Loewe additivity equation can very well be combined with existing theory on dose-effects (Chou & Talalay, 1981), causing no mathematical contradiction. The indetermination in the varying potency ratio case results of ambiguously combining dose equivalence and sham combination principle in the way Grabovsky and Tallarida suggest.

While the Loewe model does not lack uniqueness, it is nonetheless justified to doubt its validity in several cases. Among the critics of its validity in the rather generic case of varying potency ratio one finds Loewe himself (Loewe, 1953). In particular, skepticism in the case of differing maximal effects when combining a full and a partial agent has led to an increasing amount of competitive models (Greco *et al.*, 1995; van der Borght *et al.*, 2017; Twarog *et al.*, 2016; Wicha *et al.*, 2017). Experimentalists emphasize that curved rather than straight isoboles are expected (Luszczki, 2008; Geary, 2013) if dose-effect curves are not parallel, which is confirmed by examples of mechanistic models (Bosgra *et al.*, 2009). To account for these deficits, Hand introduced an alternative and more general non-interaction model (Hand, 2000), which has unfortunately been overlooked in the reception of synergy detection models. Hand suggests to construct dose-effect curves for combined agents via an ordinary differential equation (ODE) in a way that both agents contribute linearly to the instantaneous gain in effect.

The conflict about the compatibility of sham combination principle and dose equivalence has been persisting for long. In this manuscript, we show how the Hand model can be obtained as a unique limit model of the Tallarida model. In doing so, we add an argument to dissolving this conflict. We introduce effect-sensitivity curves as a visualization tool, that provides the best insight into the Hand model’s concept of additivity. This does not replace the dose-effect-visualization, but adds an alternative view, which confirms the biochemical plausibility of the Hand model in terms of the dynamical change in effect. We explore qualitative features of the Hand model that are not discussed in (Hand, 2000). Most strikingly, its isoboles are always convex, which reveals that the Hand model is systematically a more conservative prediction than Loewe. Furthermore, we demonstrate how the models by Loewe, Hand and Tallarida can be interpreted as contrasting strategies for solving to some extent an unsolvable partial differential equation characterizing the effect-surface. Finally, we compare the reference effects of different null models for a set of published dose effect data.

## 2. Mathematical background

### 2.1 Dose-effect curves and drug combinations

The dose-effect curves of the mono-therapies with drugs *A* and *B* are denoted by *f*_*A*_ and *f*_*B*_. Both curves are assumed to be strictly increasing and their inverses are denoted by 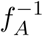 and 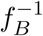. In practice, Hill functions are widely used (Motulsky & Christopoulos, 2004), i.e., dose-effect curve

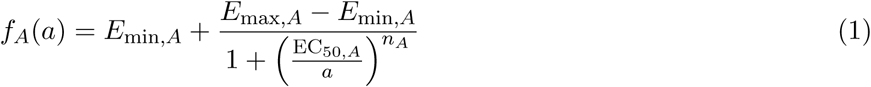

with inverse

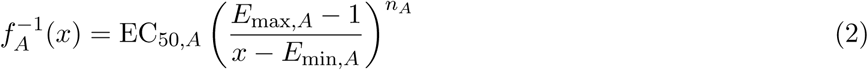

for drug *A* with dose *a*, effect level *x*, minimal effect *E*_min,*A*_, maximal effect *E*_max,*A*_, the half-max dose EC_50,*A*_ and Hill coefficient *n*_*A*_. The minimal effect *E*_min,*A*_ describes the untreated condition and is identical for different drugs, *E*_min,*A*_ = *E*_min,*B*_. The maximal effects *E*_max,*A*_ and *E*_max,*B*_ can differ and the drug with the greater maximal effect is called the full agent, whereas the other is labeled as partial agent. In general *E*_min,*A*_ is normalized to zero, and the larger of *E*_max,*A*_ and *E*_max,*B*_ is normalized to one.

The potency ratio *α*(*x*) at effect level *x* quantifies how much more potent *A* is compared to *B* in generating effect *x*, i.e.,

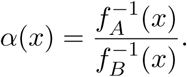

If the potency ratio is constant, both curves are identical up to rescaling the dose axis, or equivalently both curves are parallel, when plotted on a logarithmic dose axis.

We denote a combination drug of *A* and *B* at fixed ratio *λ ∈* [0, 1] by *C*_*λ*_, i.e., one unit [*c*] of the agent *C*_*λ*_ is composed as [*c*] = *λ*[*a*] + (1 *-λ*)[*b*]. Accordingly, the treatment with an amount *c* of the combination drug *C*_*λ*_ corresponds to a treatment with *a* = *λc* of drug *A* together with *b* = (1 *-λ*)*c* of drug *B*.

### 2.2 Favorable properties of a null model

Given the dose-effect curves *f*_*A*_ and *f*_*B*_ of the single agents, a null model predicts the effect as a function *E*: [0, *∞*) *×* [0, *∞*) *→* [0, *∞*),

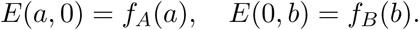

Two cuts through the two-dimensional effect-surface {(*a, b, E*(*a, b*))*| a, b ≥* 0} visualize the combined effect (Greco *et al.*, 1995): Dose-effect curves of combination drugs *C*_*λ*_ vertically cut the surface along a ray directed according to the fixed ratio *λ*, i.e., the combined curve follows

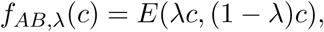

where we sometimes oppress the index *λ* for readability (Figure 1B). Isoboles are horizontal cuts at level *x*, i.e., level sets of *E*, projected onto the ([*a*], [*b*])*-*plane (Figure 1C).

A null model for drug combinations should ideally possess at least two properties to be biochemically plausible:

i. The combination of a drug with itself should neither result in synergy nor antagonism, meaning that it does not interact with itself and meets the sham combination principle. In the above notation *E* satisfies the *sham combination principle* if for all *λ ∈* [0, 1]

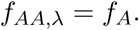

Geometrically the isoboles of the effect surface are then straight lines with unit slope.
ii. The swap of drug *A* and *B* should not change the effective results. Mathematically, this implies that E is *commutative* in *A* and *B*, i.e., for all *λ ∈* [0, 1]

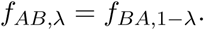

In addition, a model may satisfy an *associative property*, which allows to combine combination drugs:
iii. The combination of agents *C*_*λ*_ and *C*_*µ*_ at ratio *v* should be equivalent to directly combining *A* and *B* at suitable ratios. For [*c*_*λ*_] = *λ*[*a*] + (1 *-λ*)[*b*] and [*c*_*µ*_] = *µ*[*a*] + (1 *-µ*)[*b*], the resulting combination is *v*[*c*_*λ*_] +(1 *-v*)[*c*_*µ*_] = (*vλ* +(1 *-v*)*µ*)[*a*] +(*v*(1 *-λ*) +(1 *-v*)(1 *-µ*))[*b*]. *E* satisfies the associative principle if for any *λ, µ, v ∈* [0, 1]

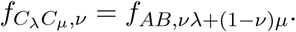

The associativity principle is stronger than commutativity, because the latter is contained in the associativity property by the particular choice *λ* = 0, *µ* = 1.

Geometrically, associativity means: *C*_*λ*_ and *C*_*µ*_ define new coordinate axes in the ([*a*], [*b*])-plane with angle smaller than 90*°*. The model prediction along these curves are taken to be the new mono-therapies while the rest of the effect surface is ignored. Bending these axes to an angle of 90*°* and applying the model to this new coordinate system or bending the null model obtained from *A* and *B* should then span the same effect surface. The associative property guarantees that the model is in this sense compatible with a change of the coordinate axes.

### 2.3 Dose-effect based null models

In this section, we introduced the null models proposed by Loewe, Tallarida and Hand. These null models are based on the dose-effect strategy and rely on the knowledge of the complete dose-effect curve.

#### 2.3.1 The Loewe model

The classical approach for mutually exclusive drug combinations is the Loewe model (Loewe, 1953; Berenbaum, 1977). As its core, the model postulates that the isoboles are straight (Loewe & Muischnek, 1926). The isobole at effect level *x* can thus be characterized as the set of all combinations (*a, b*) satisfying the Loewe additivity equation

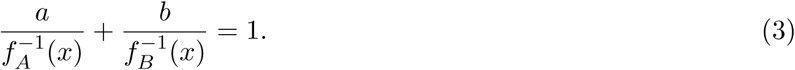

The effect *E*_*L*_(*a, b*) predicted by the Loewe model for an arbitrary combination (*a, b*) can be obtained by (numerically) solving (3) for *x*, i.e., *E*_*L*_(*a, b*) is the unique effect, that solves

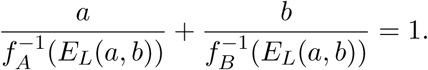

A generic point on the isobole has the form 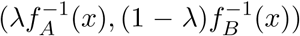 (Figure 1A). *E*_*L*_ is commutative and satisfies the sham combination principle. However, it is only defined for effects *x*, which are produced by both *f*_*A*_ and *f*_*B*_. Hence, it is not applicable for effect sizes *x ≥* min*{E*_max,*A*_, *E*_max,*B*_}.

The potency ratio *α*(*x*), which is geometrically the slope of the isobole, precisely expresses the conversion rate of equivalent doses in the Loewe model. If *A* and *B* exhibit a constant potency ratio, then all isoboles in the Loewe model are parallel. Tallarida (2006), Geary (2013) and others argue that this property is necessary to ensure validity of the Loewe model as linear isoboles are valid only if the dose-effect curves of the drugs “considered have constant relative potency”(Geary, 2013). Accordingly, the probable case of a varying potency ratio would prevent the use of the Loewe model.

#### 2.3.2 The Tallarida model

In an attempt to transfer the principles of dose equivalence and sham combination to the case of varying potency ratio, Grabovsky and Tallarida suggested to convert a dose *a* into an equivalent *B*-dose 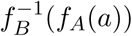 generating the same effect *f*_*A*_(*a*) (Figure 2). Adding this *B*-equivalent to *b* should predict the effect via *f*_*B*_ (Grabovsky & Tallarida, 2004), i.e.,

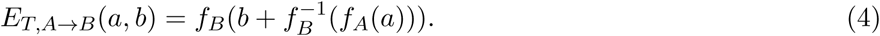

Since the model was further popularized by Tallarida (Tallarida, 2006, 2011), we refer to it as the Tallarida model. Conceptually, the Tallarida model decomposes the total effect *x* = *E*_*T,A→B*_(*a, b*) = *x*_*A*_ + *x*_*B*_ as in the following way: dose *a* of *A* individually achieves an effect *x*_*A*_ = *f*_*A*_(*a*) on its lower range of doses, while dose *b* of *B* acts on its higher dose range, contributing on top of the effect achieved by *A*, i.e.,

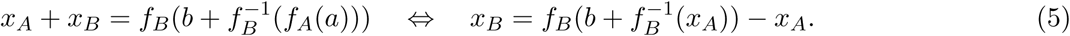

Note that the model with altered roles of *A* and *B* is

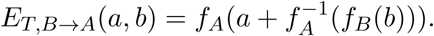

The models *E*_*T,A→B*_ and *E*_*T,B→A*_ might predict a different effect because in general

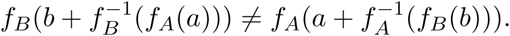

Accordingly, while the Tallarida model meets the sham combination principle, it does not fulfill the commutation property. The two model predictions *E*_*T,A→B*_(*a, b*) and *E*_*T,B→A*_(*a, b*) encompass a range of effect values. In the following, we will refer to the TallaridaLB model and the TallaridaUB model,

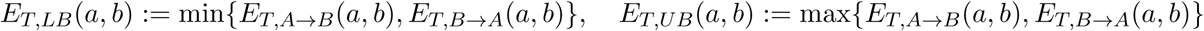

that give a lower and an upper bound to this range.

**Figure 2:**
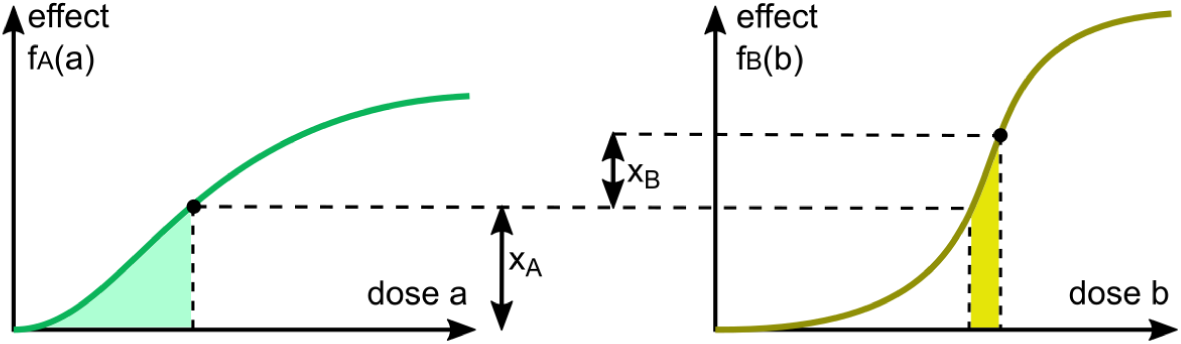
Effect decomposition in the Tallarida model. Dose *a* of *A* individually achieves an effect *x*_*A*_ = *f*_*A*_(*a*) on its lower range of doses, while dose *b* of *B* acts on its higher dose range, contributing on top of the effect achieved by *A*. The contribution of drug *B* to the effect is 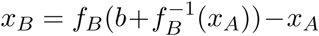.

#### 2.3.3 The Hand model

It remains the problem that the Tallarida model, introduced in 2004 by Grabovsky and Tallarida, is not consistent with changing the roles of *A* and *B*. In 2000, Hand already anticipates the ambiguity in what we call the Tallarida model. He identifies as a solution to this problem that instead of “tak[ing] the effect of [*b*] on top of whatever effect level was reached by” *a* in equation (4) one needs to “capture the fact that the drugs have their effect simultaneously” (Hand, 2000). He proposed to construct *E* by specifying the rate at which the effect is changing. Since the rate is a one dimensional quantity, it is advantageous to assign values to the entire plane of dose pairs by going along rays at fixed ratio *λ*. More precisely he suggested that the combined curve *f*_*AB,λ*_ must satisfy the differential equation

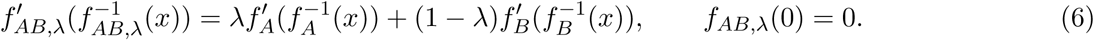

This characterizing equation states that at each effect level both agents contribute linearly to the instantaneous gain in effect of the combined curve. For an arbitrary dose pair (*a, b*), identify the ratio *λ ∈* [0, 1], such that

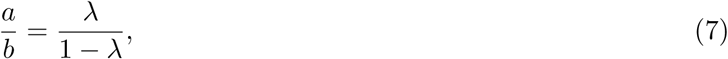

then (*a, b*) = (*λc,* (1 *-λ*)*c*) is a dose *c* of the drug mixture *C*_*λ*_. For *f*_*AB*_ solving (6) with *λ* satisfying (7), we assign

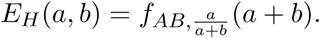

Using the inverse derivative formula, we express the differential equation (6) as

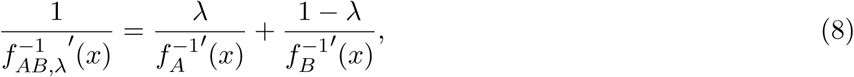

or in integral representation

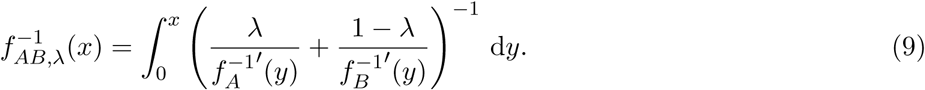

The Hand model was constructed to satisfy the sham combination principle and clearly it is commutative in *A* and *B*. Hand stresses that if an agent saturates at a lower level than the other, its derivative is set to zero above that level. This is a natural interpretation of (6) given that *x* is not in the range of either *f*_*A*_ or *f*_*B*_. The Hand model thus comprises the case of a full and a partial agent. Moreover, Hand mentions that the isoboles will be straight in the constant potency ratio case. Coinciding with the Loewe model for this subcase, it is regarded by Hand as a generalization of the Loewe model. We will demonstrate in Section 3.3, that the Loewe model is justified also for cases of varying potency ratio and thus the term “generalization” should be used with care.

### 2.4 Effect-based null models

Alternatives to the dose-effect based null models are effect-based models. These null models require only the knowledge of the effect of the single doses *a* and *b* to provide a prediction of the combined effect *E*(*a, b*). Commonly used effect-based models are the Bliss model and the HSA model. The Bliss model (Bliss, 1939) is based on a probabilistic framework, and defines the effect of a drug combination as

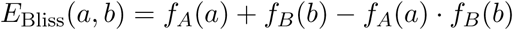

For this definition it is necessary, that the greater of the maximum values of the two dose-effect curves, *E*_max,*A*_ and *E*_max,*B*_, are normalized to 1. The HSA model, often referred to as Gaddum’s non-interaction (Berenbaum, 1989), is defined as

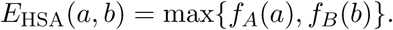

The Bliss and the HSA model do not fulfill the sham and the associative principle.

For a detailed comparison of the properties of all null models we refer to Section 3.4.1 and Table 1.

**Table 1:** Properties of null models.

|  | Loewe | Tallarida | Hand | Bliss | HSA |
| --- | --- | --- | --- | --- | --- |
| uniqueness | ✓ | — | ✓ | ✓ | ✓ |
| commutativity in $A$ and $B$ | ✓ | — | ✓ | ✓ | ✓ |
| sham combination principle | ✓ | ✓ | ✓ | — | — |
| associative property | ✓ | — | ✓ | — | — |
| defined for full and partial agent | — | ✓ | ✓ | ✓ | ✓ |

## Results

### 3.1 Mathematical analysis of the Hand model

In this study, we consider five previously published null models. Amongst them, the Hand model is the least well known and studied one. To facilitate an intuitive understanding: we will provide a unified derivation of the Tallarida and the Hand model; and we will introduce the novel concept of effect-sensitivity curves and confirm the additivity principle. Subsequently, we will study the properties of the Hand model for combinations of partial and full agents and its isoboles.

#### 3.1.1 Unified derivation of the Tallarida and the Hand model

In the literature, the Tallarida and the Hand model are presented and discussed separately (Grabovsky & Tallarida, 2004; Hand, 2000). Here, we establish a link and improve the understanding of the Tallarida and the Hand model by providing a unified derivation. By this derivation we extended the original paper’s frank justification on addition of rates.

We start from the concept of Tallarida on equivalent doses (5). However, instead of applying *a* entirely and then *b* entirely, we apply portions of both, constantly alternating (Figure 3). Formally, we split both doses *a* and *b* into *N* pieces 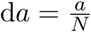 and 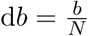, respectively. In particular it is

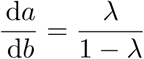

with *λ* satisfying (7). We define d*c*:= d*a* + d*b* as a small portion of the combination drug *C*_*λ*_. By alternately applying portions d*a* and d*b*, we keep track of how the effect changes. Suppose, we have constructed the dose-effect curve *f*_*AB,λ*_ of *C*_*λ*_ up to an effect level *x*_0_, induced by a mixture dose

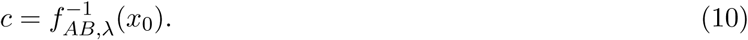

Applying d*a* elevates the effect level to 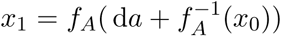 and d*b* subsequently amplifies the effect level to 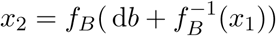. By the above procedure we obtain

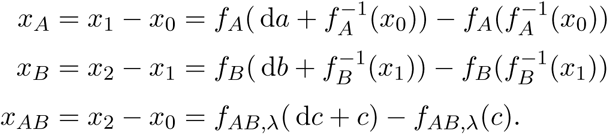

For *N* = 1, this procedure yields the Tallarida model. To study *N → ∞* we take the first-order Taylor approximation, divide by d*c* and note that d*a* = *λ* d*c,* d*b* = (1 *-λ*) d*c*. This yields

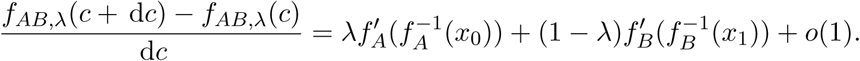

In the limit case d*c →* 0, i.e., an infinite amount of infinitesimal pieces d*c*, the difference quotient tends to the derivative on the left side in (6) and we obtain the Hand model. Note that the roles of *A* and *B* in the derivation may be switched, approaching the same differential equation in the limit. This yields a unique limit model even though each approximative model is ambiguous.

**Figure 3:**
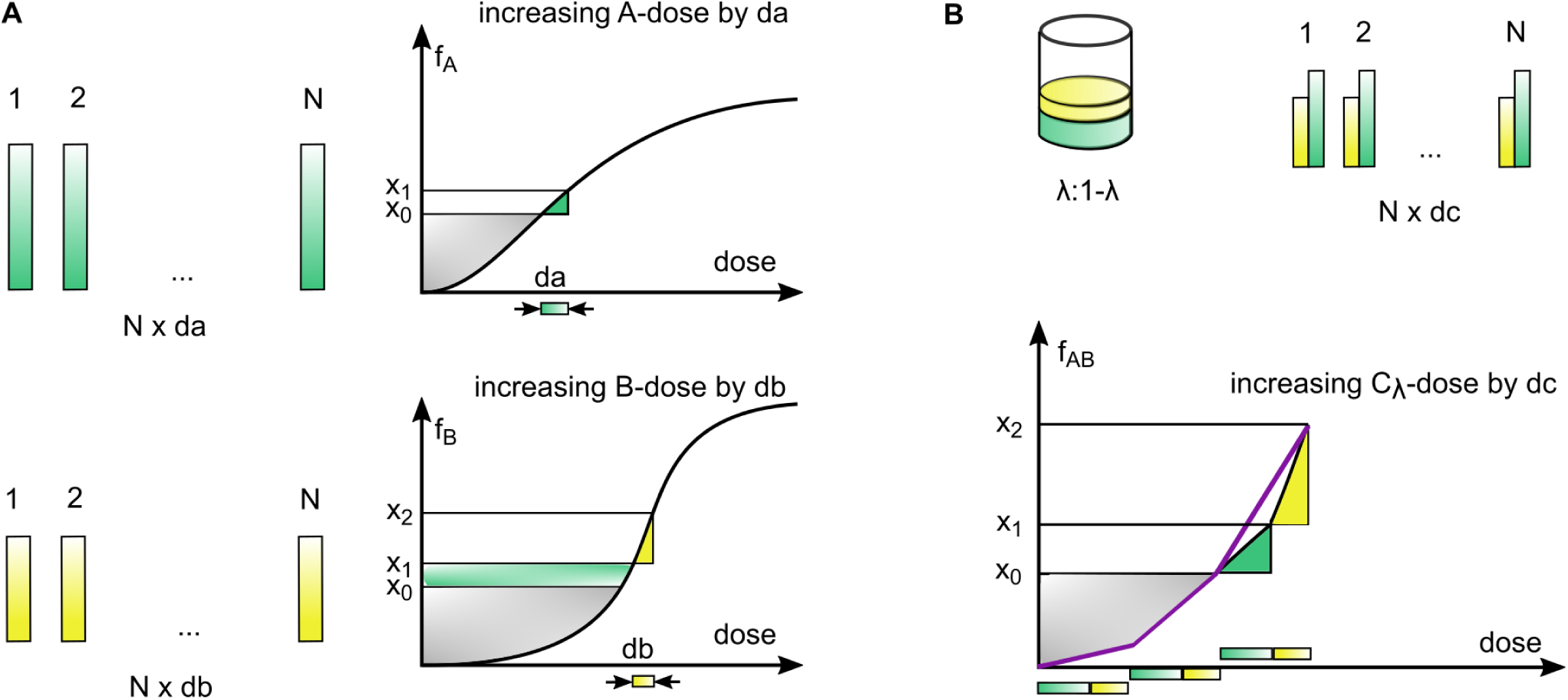
Visualization of the infinitesimal approach in the Hand model. **(A)** The doses *a* and *b* are split into *N* small pieces d*a* and d*b*, respectively. The effect level *x*_0_ is elevated by d*a* to *x*_1_, d*b* raises the effect from *x*_1_ to *x*_2_. **(B)** The combined dose-effect curve for the mixture *C*_*λ*_ is constructed by applying *N* pieces of d*c* = d*a* + d*b*. After twice applying d*c*, the effect level *x*_0_ is attained. Applying a third mixture of d*a* and d*b* successively increases the effect to *x*_1_ and finally *x*_2_.

#### 3.1.2 Effect-sensitivity curves and an additivity principle for the Hand model

In the Hand model (6) the instantaneous gain in effect depends on the individual dose-effect curves *f*_*A*_(*a*) and *f*_*B*_(*b*) via the functions 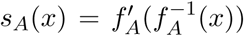 and 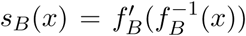, respectively. We term the derivative 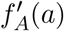, which is measured in effect per dose, its *sensitivity*. As the functions *s*_*A*_(*x*) and *s*_*B*_(*x*) describe the *sensitivity* of the dose-effect relation subject to the *effect* level, we term (*x, s*_*A*_(*x*)) and (*x, s*_*B*_(*x*)) *effect-sensitivity curves*. The dose-effect curve *f*_*A*_ is then the solution to

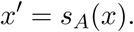

The sensitivity is precisely the right-hand side of the autonomous ODE which guides the dynamics of the dose-effect curve, evolving over the dose range instead of time. For Hill curves as in (1) the effect-sensitivity curve is explicitly given by

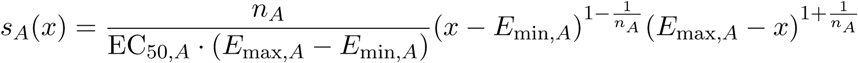

for *x ∈* [*E*_min,*A*_, *E*_max,*A*_] (Appendix 5.2.1). Under weak conditions, it is possible to retransform effect-sensitivity into dose-effect curves, rendering them an equivalent notion (Appendix 5.2.2).

The Hand model specifies the following additivity concept

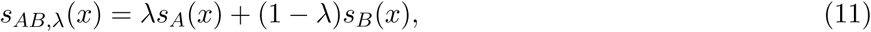

where the quantities, which are added, are the sensitivities; i.e. the effect-sensitivity curve of the combined agent is a weighted average of the single effect-sensitivity curves with weights according to the mixture ratio (Figure 4). More suggestively, we say that the Hand model summates the dynamics of the single dose-effect curves.

**Figure 4:**
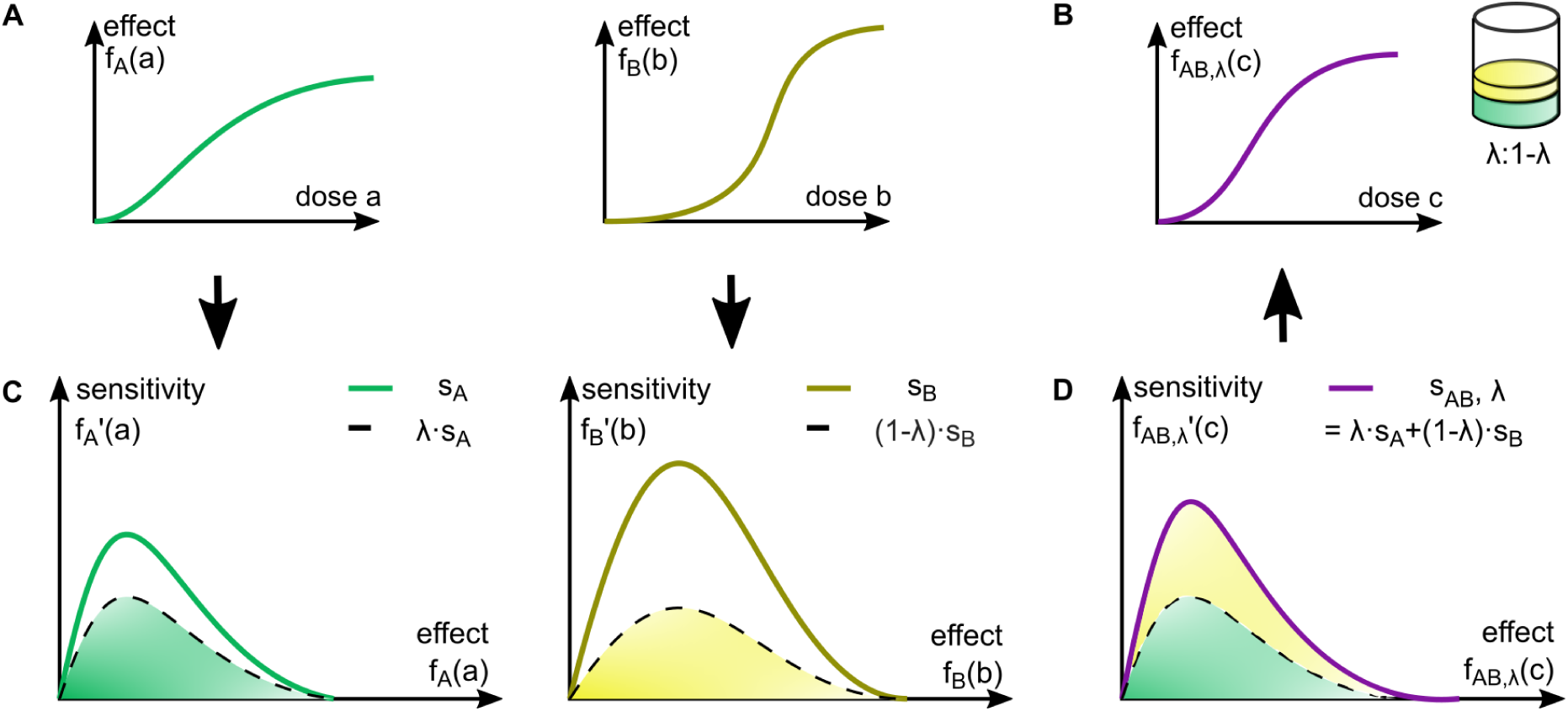
Concept of additivity according to the Hand model. **(A)** The Hand model specifies how to add the two dose-effect curves of drugs *A* and *B* to get the combined dose-effect curve at fixed dose ratio *λ* **(B)**. **(C)** *f*_*A*_ and *f*_*B*_ are transformed to effect-sensitivity curves via 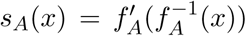, reflecting the saturation dynamics. **(D)** The effect-sensitivity curve *s*_*AB,λ*_ of the mixture *C*_*λ*_ is their weighted algebraic sum. *f*_*AB,λ*_ is obtained from *s*_*AB,λ*_ by integration.

#### 3.1.3 Geometric properties of the Hand model’s isoboles

The isoboles of the null model are the basis for assessing synergy and antagonism of drugs. While the Loewe model’s isoboles are easily graphically obtained and analytically given in closed form as soon as 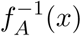 and 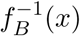 are known, the Hand model’s isoboles are nontrivial and might often have to be computed numerically. However, we could show that the Hand isoboles share a characteristic geometric feature: they are always convex (Appendix 5.4). As a consequence, they systematically diminish the region of synergy compared to the Loewe model.

### 3.2 Combination drugs for partial and full agent

In practice the maximal effect of drug *A* might be smaller than the maximal effect of drug *B*, meaning that *A* acts as the partial agent and *B* as the full agent. Since the construction in the classical Loewe model requires effect levels that are attained by both agents (Loewe, 1953), this case has motivated to a large extent the search for alternative models (Greco *et al.*, 1995; van der Borght *et al.*, 2017; Twarog *et al.*, 2016; Wicha *et al.*, 2017). In the following we assume *E*_max,*A*_ *< E*_max,*B*_.

#### 3.2.1 Loewe model for partial and full agent

The Loewe model assigns effect values to all dose pairs (*a, b*) with 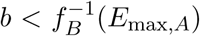 (Figure 5A). In fact, the slope of the isoboles approaches 0 as the effect level approaches *E*_max,*A*_. This phenomenon arises because the Hill curve *f*_*A*_ saturates asymptotically. Hence the strip in the ([*a*], [*b*])-plane that is bounded by the horizontal line 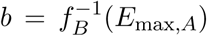 is covered entirely by Loewe isoboles. The horizontal limit isobole 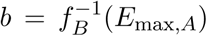 coincides with the HSA model’s isobole at level *E*_max,*A*_, which suggests to extend the Loewe model by the HSA model.

**Figure 5:**
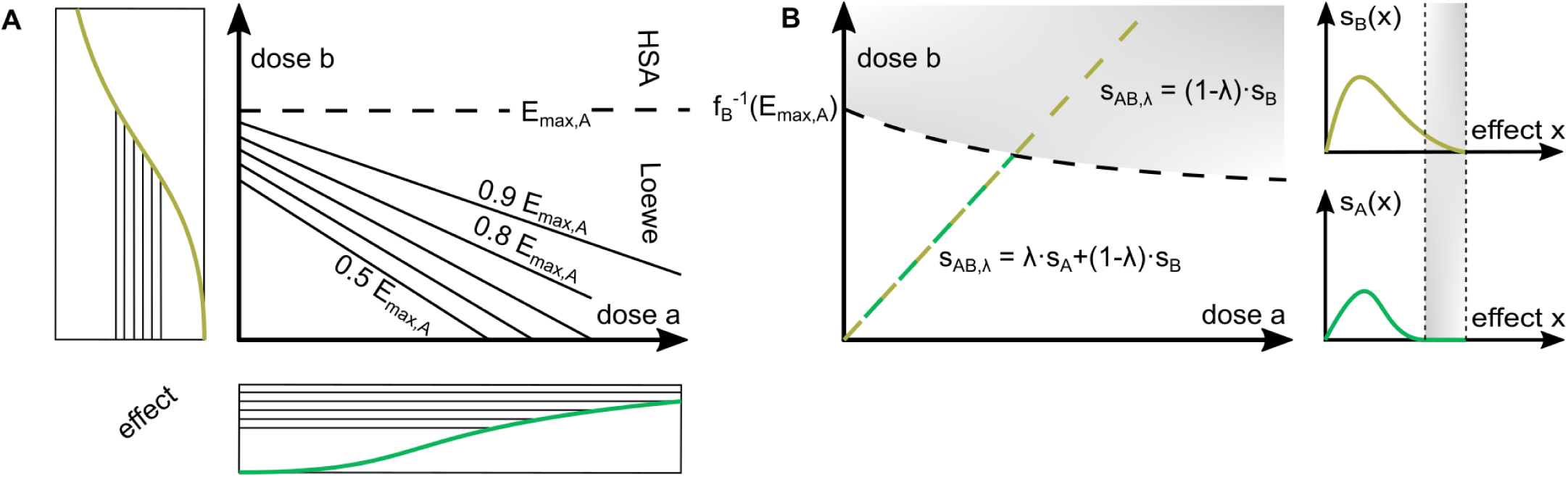
Model predictions for combination drugs of a partial agent *A* **and full agent** *B*. The isobolograms for the **(A)** Loewe model and **(B)** Hand model. **(A)** The Loewe model assigns reference effects only to dose pairs below the horizontal limit isobole at effect value *E*_max,*A*_, beyond which it can be extended via the HSA model. **(B)** Above the partial agent A’s maximal effect value its sensitivity is set to 0. As soon as the combined curve passes the curved limit isobole at effect level *E*_max,*A*_ it is driven by the full agent alone.

#### 3.2.2 Hand model for partial and full agent

As shown in (Hand, 2000), the Hand model comprises the case of a partial agent *A* and a full agent *B*. In terms of sensitivities, the effect-sensitivity curve *s*_*A*_ is extended by constant zero on the effect domain [*E*_max,*A*_, *E*_max,*B*_) (Figure 5B), i.e. the sensitivity of the combined curve is

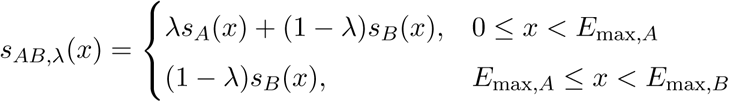

If *c*^*\**^(*λ*) denotes the dose of mixture *C*_*λ*_ at which *E*_max,A_ is attained, then

- for *c ≤ c*^*\**^(*λ*), the ODE guiding the evolution of *f*_*AB,λ*_ is (6).
- For *c > c*^*\**^(*λ*), the evolution simplifies to the initial value problem

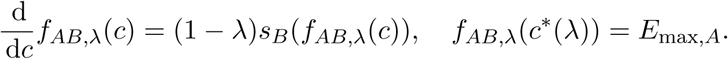

The solution for *c > c*^*\**^(*λ*) is 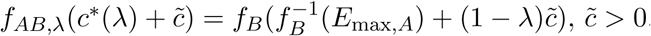, hence,

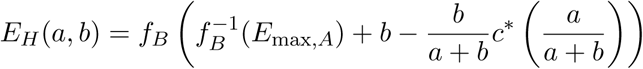

for all 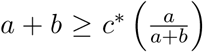. Accordingly, for effect sizes greater than *E*_max,*A*_ only B has an effect and the dose-effect curve for *C*_*λ*_ in this regime corresponds to the scaled and shifted dose-effect curve *f*_*B*_. In contrast to the Tallarida model, the effect surface does not copy the dose-effect curve *f*_*B*_ in the direction parallel to the *B*-axis.

Using the integral representation, *c*^*\**^(*λ*) is explicitly given by

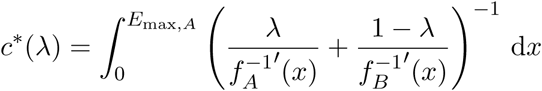

Decomposing the combined dose *c*^*\**^(*λ*) into its contributions *a*^*\**^(*λ*) = *λc*^*\**^(*λ*) and *b*^*\**^(*λ*) = (1 *-λ*)*c*^*\**^(*λ*), the limit isobole is parametrized by *λ 1→* (*a*^*\**^(*λ*), *b*^*\**^(*λ*)). Its *B*-component

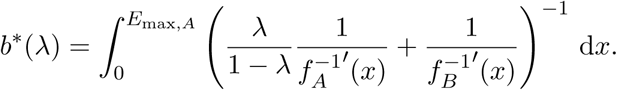

is a monotone continuous function with lim_*λ→*1_ *b*^*\**^(*λ*) = 0.

Let us contrast this finding with the Loewe model. Assuming Hill curves, which saturate asymptotically, as dose-effect-relations for both models, the Loewe model ignores the amount of *A* at the critical effect size *E*_max,*A*_, while in the Hand model a higher amount of *A*, and thus a larger *λ*, reduces the amount of *b*^*\**^(*λ*) necessary to generate the effect *E*_max,A_.

### 3.3 Common framework for the Loewe, the Tallarida and the Hand model

The dose-effect based null models presented above pursue different approaches in constructing the effect surface *E*(*a, b*). While the Loewe model imposes a condition on the isoboles’ shape, the Tallarida model elevates the dose-effect curves to the respective initiating level of the other curve. Consequently it constructs the effect surface *E*(*a, b*) by copying and attaching the existing curves of the single agents. The Hand model specifies the evolution of effect surface *E*(*a, b*) for increasing amount of mixture dose at fixed ratio, hence constructs the surface along rays. While the models seem rather different, we could derive a novel common framework for all three, based on the concept of effect-sensitivity curves introduced in Section 3.1.2.

Assuming an underlying mechanistic dynamical model for the effect of the individual drugs, it is plausible to request that the change in effect should only depend on the current effect level. If nothing but the knowledge about the single dose-effect curves is available, it is plausible that the single dynamics predict the dynamics of the combined effect. More precisely, we expect the sensitivities of *f*_*A*_ and *f*_*B*_ to translate changes of *A*-and *B*-doses respectively into changes in effect, i.e.,

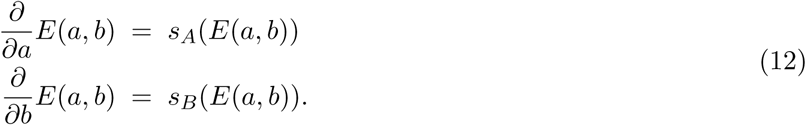

For given dose-effect curves *f*_*A*_ and *f*_*B*_, and the derived effect-sensitivity curves *s*_*A*_ and *s*_*B*_, such a function *E* does in general not exist. However, if we request the equations to hold to some extent, we recover the above null models.

The **Loewe model** is recovered, if we request (12) to hold along the isoboles (Figure 6A): Let *γ* = (*γ*_*A*_, *γ*_*B*_): [0, 1] *→* [0, *∞*) *×* [0, *∞*) parametrize the isobole at effect level *x*, then

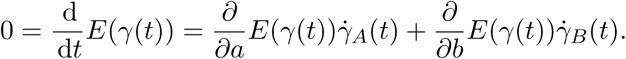

**Figure 6:**
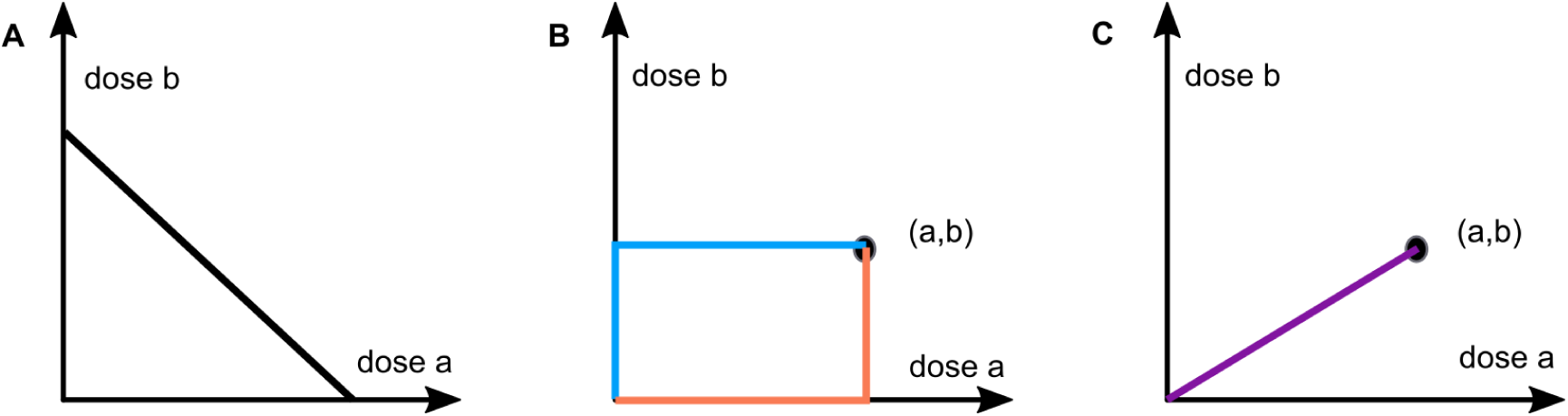
Paths for the three null models. The three null models solve equation (12) to some extent along different paths. **(A)** The Loewe model solves (12) along isoboles. This requirement forces the isoboles to be linear. **(B)** The Tallarida model solves (12) along rectangular paths. **(C)** The Hand model solves (12) along rays.

Using *E*(*γ*(*t*)) *= x*, we obtain

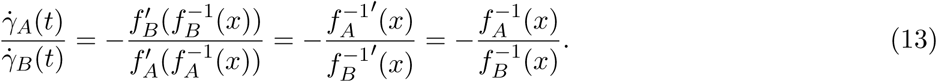

The last equality holds because 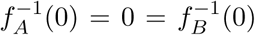. The calculation (13) shows that the ratio 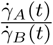 in fact not depend on *t*, but is constant for all *t*. Consequently, *γ* parametrizes a straight line with the slope given precisely by the potency ratio *α*(*x*). It is important to notice, that we did not request the isoboles to be straight lines in the first place, however, satisfying (12) forces them to be linear.

The **Tallarida model** *E*_*T,A→B*_ is recovered by following a rectangular path from (0, 0) to (*a, b*) (Figure 6B).

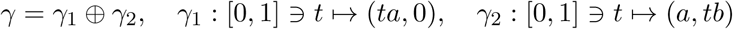

where ⊕ concatenates the paths, i.e. for *t ∈* [0, 2]

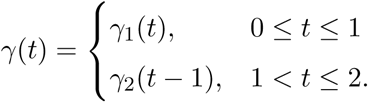

Then along *γ*_1_ equation (12) integrates to *E*_*T,A→B*_(*γ*_1_(*t*)) = *f*_*A*_(*ta*) and along *γ*_2_ equation (12) with initial condition *E*_*T,A→B*_(*γ*_2_(0)) = *E*_*T,A→B*_(*γ*_1_(1)) = *f*_*A*_(*a*) integrates to 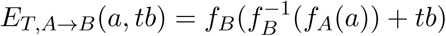. Analogously, the model *E*_*T,B→A*_ is obtained by the rectangular path

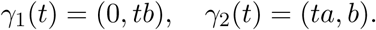

The **Hand model** is recovered if we request (12) to hold along combined curves *γ*(*c*) = (*λc,* (1 *-λ*)*c*) (Figure 6C). In this case we compute by the chain rule

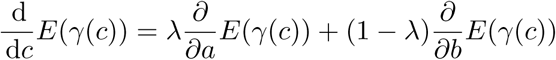

and set 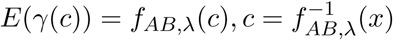.

In the constant potency ratio case, (12) admits a solution and all three models coincide. Indeed, for 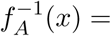 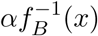 the solution *E* is given by

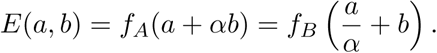

The representation of the Loewe, the Tallarida and the Hand model as alternative paths through an underlying null model (12) provides a novel perspective. It highlights the relation of the existing dose-effect based null model and the importance of the effect-sensitivity curves. Furthermore, this common framework might facilitate the basis for the development of alternative models with novel properties.

### Comparison of the mathematical properties of the null models

The null models considered in this study possess different (combinations) of properties. In this section, we collect the available information, present results on the associative principle and outline the implications of our findings. In addition, we study the reference effect values calculated by different null models and prove rigorous orderings.

#### 3.4.1 Properties of the Loewe, the Tallarida and the Hand models

Given a general framework for all three null models, we studied their properties in further detail. A summary is provided in Table 1. As outlined before,

- the Loewe, the Tallarida and the Hand model meet the sham combination principle (Section 2.3), which is not the case for the Bliss and the HSA model; and
- the Loewe and the Hand model meet the commutation principle.

Indeed, we could also prove here that

- the Loewe and the Hand model fulfill the strong property of associativity (Appendix 5.3.1).

For drugs with constant potency ratios, the Loewe and Hand model coincide. Furthermore, the Tallarida and the Hand are applicable to a combination of full and partial agents.

Our analysis reveals that the Hand model fulfills all the previously defined favorable properties (see Section 2.2 and (Hand, 2000)), while all other models only fulfill some of them. As the favorable properties are biochemically motivated, this implies that the Hand model is biochemically most plausible among the considered models.

#### 3.4.2 Systematic inequalities

The assessment of drug synergy or antagonism for a single dose of a combination drug is equivalent to the question: On which side of the isobole corresponding to the observed effect does the measured dose pair lie? Accordingly, the definition of drug synergy or antagonism depends directly on the isobole, which potentially differ substantially between the null models. For the example shown in Figure 7, the HSA, the Bliss, and the TallaridaLB model classify a dose pair to be synergistic, while the Loewe, the TallaridaUB, and the Hand model classify it as antagonistic. Note that a null model which predicts effects yields and isobole which is further away from the origin.

**Figure 7:**
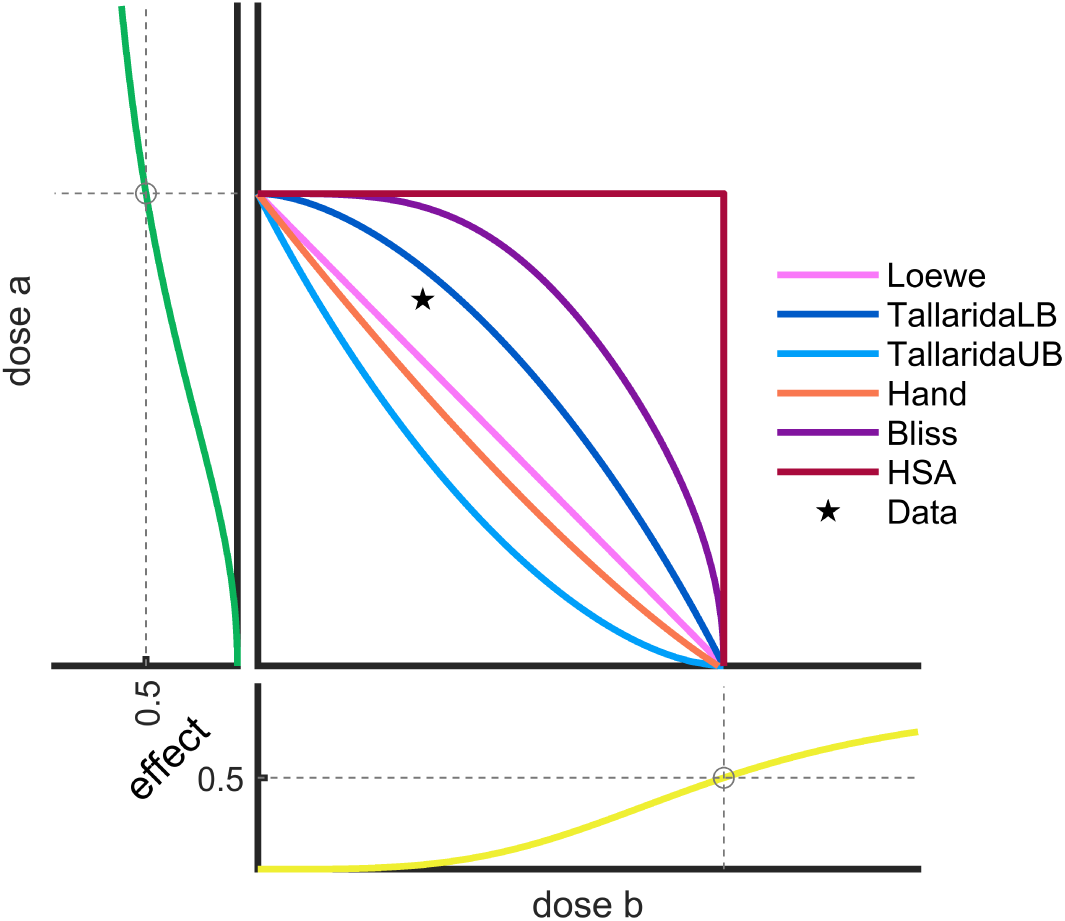
Assessment of synergy and antagonism using different null models. Dose-effect curves and isobologram at *EC*_50_ for different null models. A measured effect of 0.5 for the dose pair (10^−3^, 1.5) is considered as synergistic by the Bliss, the HSA and the TallaridaLB model, while it is antagonistic according to the Loewe, the Hand and the TallaridaUB model.

Mathematically, an important implication of the convexity of the Hand model’s isoboles that we showed earlier is that the reference effect provided by the Hand model upper bounds the reference effect provided by the Loewe model. For a specific dose pair (*a, b*), the Loewe model provides a reference effect *x* = *E*_*L*_(*a, b*). If the dose pair is expressed in terms of a quantity *c* of the mixture drug with ratio 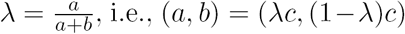, then the convexity of the Hand model’s *x*-isobole implies that 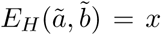 for 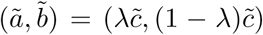 with some 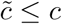. As the combined curve *f*_*λ,*Hand_(*c*) is an increasing function in *c*, we conclude

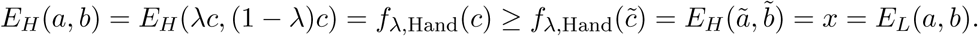

In addition, (i) TallaridaLB is by definition lower than TallaridaUB and (ii) the HSA model predicts systematically lower effects than all other null models, for instance

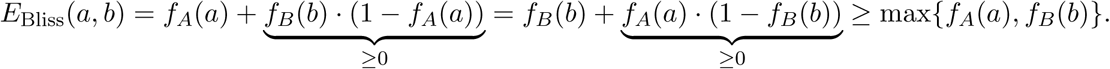

In summary, following systematic inequalities hold for the reference effect values obtained by the different null models for any combination drug *C*_*λ*_ and combination drug concentration *c*:

- HSA *<* Loewe *<* Hand
- HSA *<* Bliss
- HSA *<* TallaridaLB *<* TallaridaUB

### Comparison of null models using experimental data

To assess the characteristic features of the different null models in practice, we considered the drug screening data collected by O’Neil *et al.* (2016). The dataset reports the effects of 39 cancer cell lines to 38 drug individual drugs and 583 drug combinations of these drugs. Each combination was measured on a 4 by 4 dosing regime. The proliferation compared to the untreated condition is taken as the readout of the effect.

For the given data, we compared the reference effects of the Loewe, the Tallarida and the Hand model, as well as the Bliss model and the HSA model. As the Tallarida model does not provide a unique solution (see Section 2.3.2), we evaluated both cases. The Tallarida models predicting lower and higher effects for a particular drug combination are as before denoted as the TallaridaLB and the TallaridaUB model, respectively. For the Loewe, the Tallarida and the Hand model, Hill curves and a zero-effect curves were fitted to the measurement data for the single drug treatments. The Bayesian Information Criterion (BIC) (Schwarz, 1978) was used to select the more suited model.

For a comparison of the null models, we considered the predictions of all tested dose pairs of all tested drug combinations. For the cell line A375, the predictions are visualized in Figure 8. Besides the systematic inequalities which have been proven in Section 3.4.2, we found:

**Figure 8:**
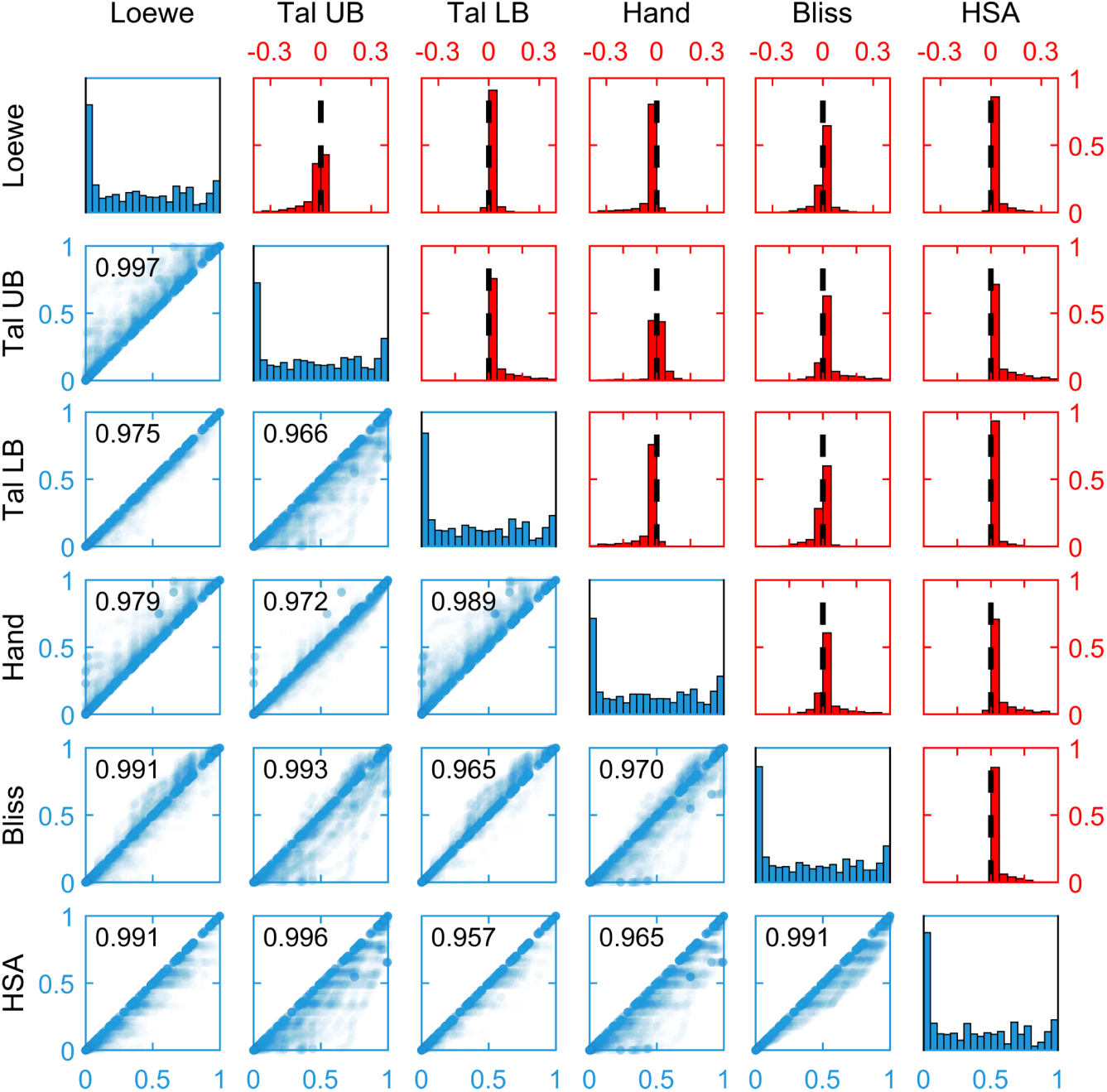
Distributions of reference effects: Below the diagonal a scatter plot of the null model reference effects for all drug doses tested for cell line A375 is shown. On the diagonal the distribution of the reference effects are shown. Above the diagonal a histogram of the differences of the references are given. Zero is marked by a dashed line.

- Reference effects for different null models were highly correlated.
- The TallaridaLB model tended to lower bound the references of all other models besides HSA, while the TallaridaUB model tended to upper bound the references of all other models besides the Hand model.
- The TallaridaLB and the TallaridaUB are potentially quite different. This can lead to problems if the researcher is not aware of the ambiguity of the Tallarida model. The analysis then gives a random choice between the lower and upper prediction and no consistent and meaningful result will be obtained.
- The Hand model provided relatively high reference effects, as suggested by theory (Section 3.4.2), and lied close to the TallaridaUB model.
- The Bliss model neither systematically provided lower nor higher reference effects than the Loewe, the TallaridaUB and the Hand model (Figure 8).

## 4 Discussion

The Loewe model has persisted for long as the first choice to model additive behavior and its validity is in general accepted in the case of constant potency ratio. Skepticism about the applicability in the case of varying potency ratio case has put forth a large variety of null models. For a full and partial agent the Loewe model is in fact incapable of providing a reference effect in a certain dose range, because the reference relies on a shared effect level of both agents. If both drugs reach the same maximal effect, a Loewe assignment is possible, even if the shapes of the saturation curves are different. This case required further arguments why to doubt the Loewe model’s validity. In this manuscript, we showed that the skepticism can be perpetuated only to some extent, see below.

It has been observed that the principle of dose equivalence and the sham combination principle very well characterize the Loewe model in the constant potency ratio case (Berenbaum, 1977; Grabovsky & Tallarida, 2004; Geary, 2013). They seem promising if one attempts to modify the Loewe reference. The coupling of both principles has served as sufficient condition (“Its [the Loewe equation’s] derivation took no account of the shapes of the dose-effect curves” (Berenbaum, 1977)), necessary condition (“The Loewe additivity rests on” (Foucquier & Guedj, 2015)) and proof of non-validity (Grabovsky & Tallarida, 2004; Geary, 2013) to the Loewe model all alike. We argued that the two principles are compatible only if the two agents have a constant potency ratio. The approach suggested by Tallarida converts doses of different drugs by a rule of three, choosing as intermediate linking quantity an absolute effect level. Two doses which attain this effect level are considered equivalent. This notion of equivalence fails as soon as it is used for adding doses. Since one dose is applied on top of the other, it must thus be interpreted as change in dose of the former. This asymmetry in the roles as initial dose and change in dose leads to the observed non-commutativity. If one tries to resolve the ambiguity by choosing as intermediate linking quantity a change in effect rather than an effect level, one merely shifts the ambiguity. A change in dose cannot be assigned a change in effect uniquely, unless the dose-effect follow a linear relation which is rarely the case.

The above ambiguities have been addressed multiple times in the literature, but Hand’s substantial suggestion of a model that resolves them has been most ignored up to now. The Hand model interprets the change in effect in an instantaneous sense and allows the instantaneous change to depend on the current effect level, when constructing the combined curve. This dependency of the instantaneous change on the current quantity level is the common motive of mechanistic approaches and is appreciated in biochemical modeling because its specification is of a local instead of a global nature. The local nature of the Hand model’s additivity concept makes it biochemically plausible even though its black box approach does not incorporate knowledge on explicit molecular reactions. We showed how these instantaneous changes are best visualized when plotted subject to the effect. The resulting effect-sensitivity curves are added in the Hand model. Here, it is crucial to choose the effect as x-axis. The mechanism of the Hand model can be applied in any case in which two increasing curves must be added. Instead of summating them as functions, their dynamics are added. This not only makes the black box ansatz biochemically plausible but contributes to its versatile usage. If two dose-effect-curves exhibit a piecewise positive derivative and share a zero level, this model is applicable. For instance, this holds in the case, in which ideal data points are linearly interpolated. The extension of the Hand model to multi-drug combinations with more than two components is straightforward, as is the case with the Loewe model.

What model should be preferred? All three models satisfy the sham combination principle, which is for instance not the case in Fisher’s dosage orthogonality or the Bliss model. The Tallarida model can be used if an asymmetry is justified, e.g., by a temporal delay. The Tallarida model can also be used to identify not only a curve but an area of zero-interaction. We have demonstrated by the common framework for all three models, that there is actually no reason to dismiss Loewe in the varying potency ratio case. The condition, which is provided in form of a partial differential equation and which is biochemically plausible is just as well satisfied to a limited extent by the Loewe model as it is by the Hand model. We suggest to rehabilitate the Loewe model for the case of shared maximal effect, but differing Hill coefficient. However, note that the Hand model can predict an effect for the combination of a full and a partial agent without restrictions, whereas the Loewe model fails to do so for a range of dose pairs. Moreover, by proving the convexity of the isoboles we demonstrated that the Hand model guarantees a more conservative prediction of zero-interaction compared to Loewe.

In summary, the interest in combination drug therapy is steadily increasing and an assessment requires suitable null models. Here, we have compared several established models and have shown that the Hand model is of most versatile use, for its unique assignment, mathematical plausibility, simplicity, conservative prediction and biochemical interpretability.

## Author contributions

C.L. and J.H. conceived the project. M.S. carried out the mathematical analysis of the null models. J.V. implemented the null models and analyzed the differences for the considered dataset. All authors contributed to the preparation of the article. All authors read and approved the final manuscript.

## Competing Interests

The authors declare that they have no competing interests that might be perceived to influence the results and/or discussion reported in this paper.

## 5 Appendix

### 5.1 Derivation of the ODE governing the combined dose-effect curve of the Hand model

According to the Hand model, the dose-effect *f*_*AB,λ*_ for the combined agent *C*_*λ*_ satisfies the ODE

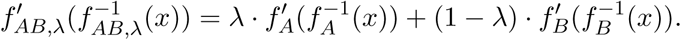

This ODE for the combined dose-effect curve *f*_*AB,λ*_ at fixed dose ratio *λ* can be derived using a Taylor expansion up to first order.

Denote by 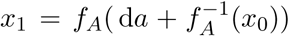 and 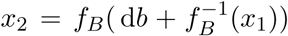 the intermediate and final effect level after applying d*a*and d*b*, respectively. Then the gains in effect due to *A, B* and their combination are

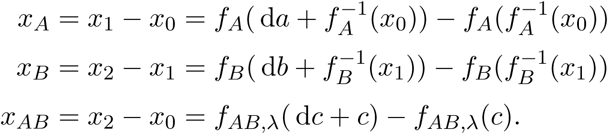

Expand

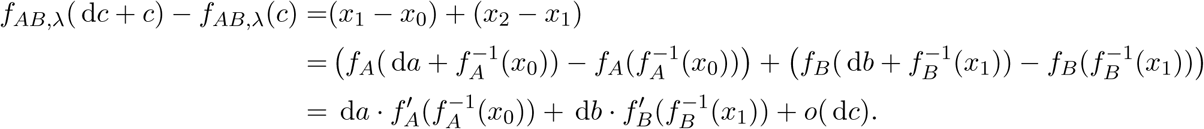

Dividing by d*c* yields

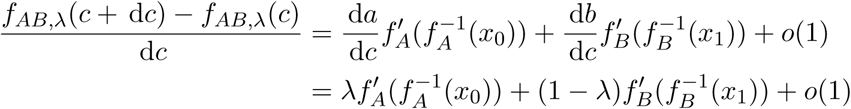

If we let tend d*c →* 0, also *x*_1_ *→ x*_0_, the difference quotient approaches

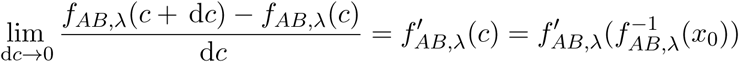

and we get the above ODE.

### 5.2 Effect-sensitivity curves

#### 5.2.1 Derivation of the effect-sensitivity formula for Hill curves

The dose-effect behavior is often modeled by Hill curves. For Hill curves of the form

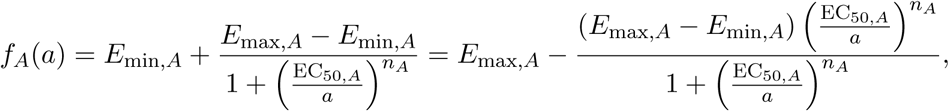

the derivative is

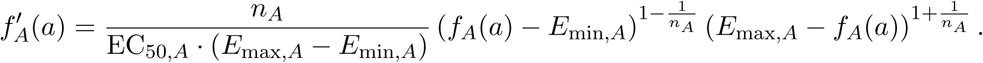

Thus, for the sensitivity we find

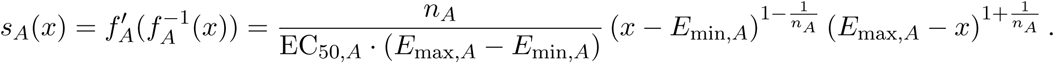

#### 5.2.2 Conversion of dose-effect and effect-sensitivity curves

Starting with a strictly increasing, piecewise differentiable dose-effect function *f*_*A*_, its inverse exists and its sensitivity is given by 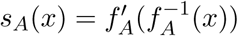. For a given positive piecewise continuous effect-sensitivity function *s*_*A*_, the inverse 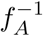 is obtained by

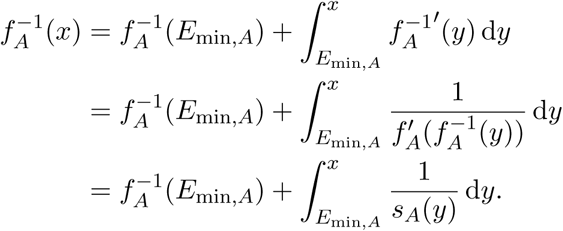

Since by this calculation 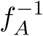 is a strictly increasing function, it is injective and its inverse *f*_*A*_ exists. Alternatively, *f*_*A*_ is obtained as the unique strictly increasing solution of the autonomous ODE *x*^*l*^ = *s*_*A*_(*x*), *x*(0) = *E*min,*A*.

In order for the above calculation to be valid, we must assume that if lim_*x→E*min,*A*_ *s*_*A*_(*x*) = 0, the integral 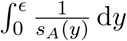 is finite. For the class of *s*_*A*_ as given by the above formula for Hill curves, this assumption is satisfied, because 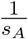 behaves like (*x - E*_min,*A*_)^*-γ*^ near *E*_min,*A*_ with 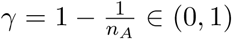 and thus integrates to a finite value. Note, that in the limit case *n*_*A*_ = *∞,* EC_50,*A*_ = *∞*, such that 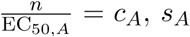, behaves like the logistic equation *s*_*A*_(*x*) = *c*_*A*_ (*x - E*_min,*A*_) (*E*_max,*A*_ *- x*) and the solution *x*(*a*) of *x*^*l*^ = *s*_*A*_(*x*) approaches *E*_min,*A*_ only in the limit *a → -∞*, not in a finite amount of dose.

Numerically the integral 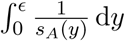 or equivalently the ODE *x*^*l*^ = *s*_*A*_(*x*), *x*(0) = *E*_min,*A*_ must be treated with care whenever *s*_*A*_(*E*_min,*A*_) = 0 because it allows the unfavored constant solution *x ≡ E*_min,*A*_ as well as solutions that are initially constant and exit *E*_min,*A*_ at an arbitrary dose value.

In the numerical implementation we solved this problem by setting the ODEś initial value to *x*_0_ = *ε* = 1e-7. Details can be found in the Appendix 5.8.

### 5.3 Model properties

#### 5.3.1 Proof of the associative property for the Loewe and the Hand model

The Loewe model: In terms of the combined curve *f*_*AB,λ*_, we write the Loewe isobole equation as

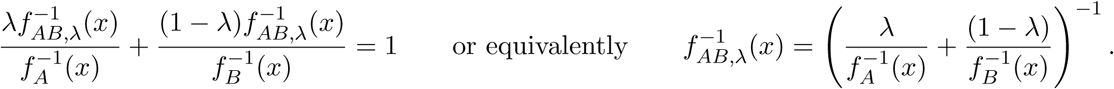

Using analogous equations for *f*_*AB,µ*_ and *f*_*Cλ*_*Cµ,v*, we can calculate

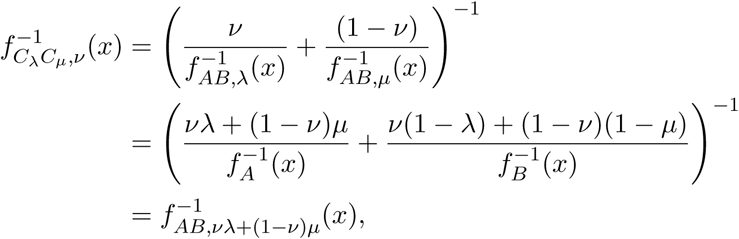

consequently 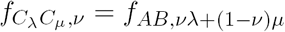.

(ii) The Hand model: The combined agents *C*_*λ*_ and *C*_*µ*_ satisfy (8)

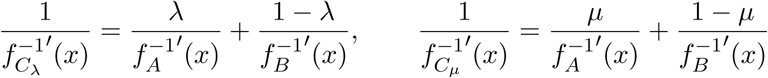

For arbitrary *v ∈* [0, 1], the grand child agent satisfies

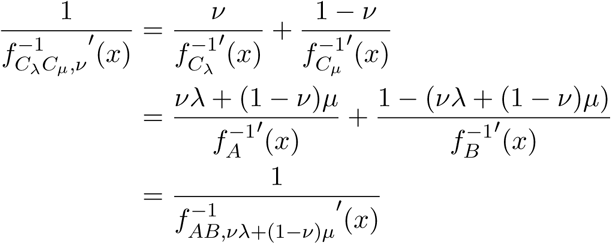

provided 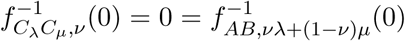 the grand child agent formed of the child agents *C*_*λ*_ and *C*_*µ*_ can indeed be formed by the parent agents *A* and *B* at appropriate ratio *vλ* + (1 *-v*)*µ*.

#### 5.3.2 Disproof of the associativity property for the Tallarida, the Bliss and the HSA model

i. The Tallarida model does not satisfy the associativity property. By the choice *λ* = 0, *µ* = 1, the associativity property implies commutativity, which the Tallarida model violates.
ii. For the Bliss model, the case 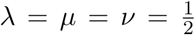, *A* = *B* shows that the associative property is violated because

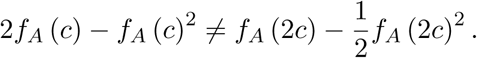
iii. The HSA model is characterized by isoboles that form a rectangle with the dose axes. If the newly allocated coordinate axes are bent, the isobole will be reshaped to an angle of more than 90*°*. The characteristic property is then lost. Moreover, the isobole suggested by applying the HSA model on *C*_*λ*_ and *C*_*µ*_ encloses the original isobole, resulting in a smaller prediction value. This geometric interpretation of the associative property is displayed in the formal definition as well: From

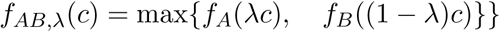

we conclude that

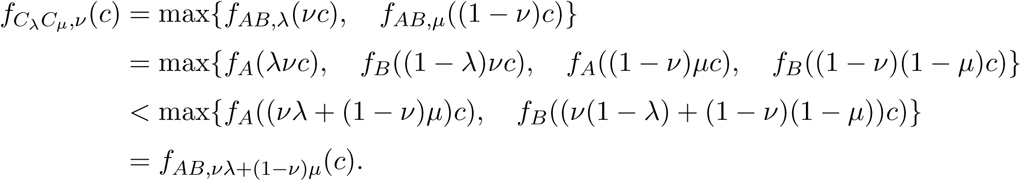

#### 5.3.3 Proof of congruency of the Hand and the Loewe model assuming constant potency ratio

Let *α* be the constant potency ratio, i.e., for any effect level *x*, 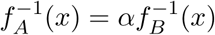, it holds for the derivatives:

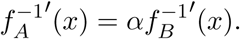

Then by (8) it follows for the combined curve *f*_*AB,λ*_ of child agent *C*_*λ*_

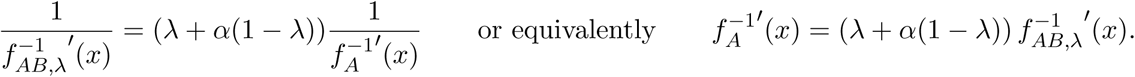

Provided that 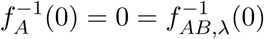, we get the relation

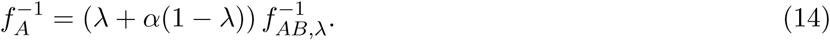

Consequently *f*_*AB,λ*_, *f*_*A*_ and *f*_*B*_ are pairwise in constant potency relation. Let *x* be a given effect level. Now, we prove that the dose pair (*a, b*) lies on the straight line connecting 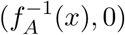 and 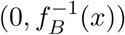, if and only if *f*_*AB,λ*_, *f*_*A*_(a + b) 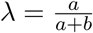:

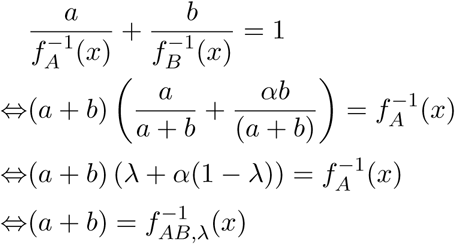

#### 5.3.4 Proof of the commutativity for the Hand, the Loewe, the Bliss and the HSA model

i. The Hand model is commutative in *A* and *B*. *f*_*AB,λ*_ = *f*_*BA,*1*-λ*_ because both satisfy the same ODE. Hence switching the roles of *A* and *B* along with their weights does not alter the combined dose-effect curve.
ii. The formulas for effects *E*_Bliss_ and *E*_HSA_ are symmetric in *A* and *B*.
iii. The Loewe isobole equation is symmetric in *A* and *B*. The isoboles determine the effect surface uniquely.

#### 5.3.5 Disproof of the commutativity for the Tallarida model

The proof for the asymmetry in the Tallarida model is given in (Lorenzo & Sánchez-Marin, 2006).

#### 5.3.6 Proof of the sham combination principle for the Hand, the Loewe and the Tallarida model

If *A* = *B*, then *A* and *B* have constant potency ratio *α* = 1. In this case the Loewe, the Hand and the Tallarida models coincide, i.e.

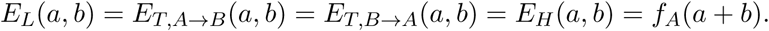

Then

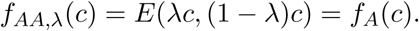

#### 5.3.7 Disproof of the sham combination principle for the Bliss and the HSA model

The sham combination principle for the Bliss model is addressed in (Foucquier & Guedj, 2015). The HSA model predicts

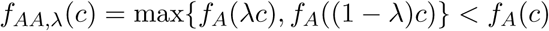

if *λ ∈* (0, 1) and *f*_*A*_ is strictly increasing.

### 5.4 Proof of the isoboles’ convexity in the Hand model

The isoboles obtained from the Hand model are convex. They are strictly convex if and only if *f*_*A*_ and *f*_*B*_ exhibit a varying potency ratio.

We prove this property of the Hand model using (i) the integral representation (9) and (ii) the associativity property:

(i) Fix an effect level *x*, which is in the target domain of *f*_*A*_ and of *f*_*B*_. Then for any *λ ∈* (0, 1), it holds with 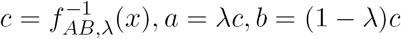, that

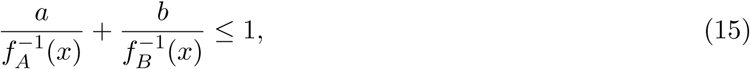

i.e., the pair (*a, b*) predicted by the Hand model to generate *E*_*H*_ (*a, b*) = *x* lies below the Loewe straight isobole.

(ii) To show convexity, apply the same argument with two child agents *C*_*λ*_, *C*_*µ*_ instead of *A, B* and corresponding points

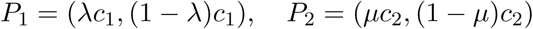

on the isobole at level *x*. By the associativity property, we can conclude that for any *σ ∈* [*λ, µ*] the point (*σc,* (1 *-σ*)*c*) on the Hand isobole at effect level *x* lies below the line segment through *P*_1_ and *P*_2_, completing the proof of the convexity property.

Now that the plan was established we carry out the steps.

(ii) We proceed by first proving (15). For readability set 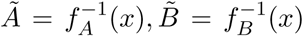. Factoring out *c*, and multiplying by *ABλ*^−1^(1 *-λ*)^−1^, (15) is equivalent to

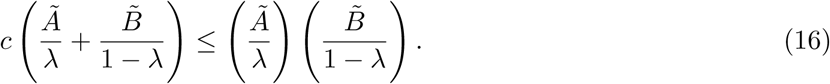

Integrating the rates 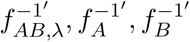, we get the following representations

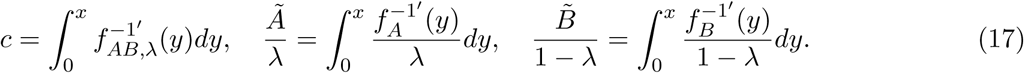

Define the second and third integrand as *h*_*A*_(*y*) and *h*_*B*_(*y*), respectively. Using the integral representation (9) of the ODE and the identity 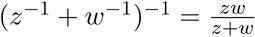, which holds for any two positive real numbers:

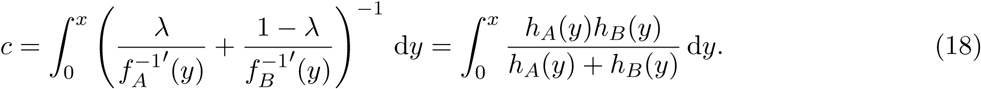

Substituting (17) and (18), (16) is equivalent to proving:

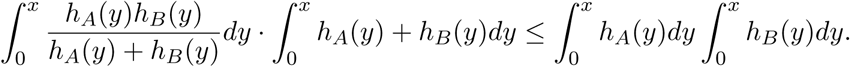

In order to establish this integral inequality, we use Cauchy-Schwarz

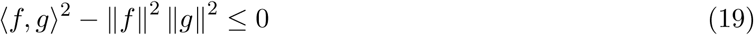

for the scalar product

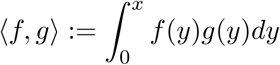

where we choose 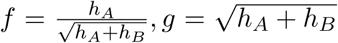. For readability we omit the integral bounds 0, *x* and the integration variable *y* when calculating

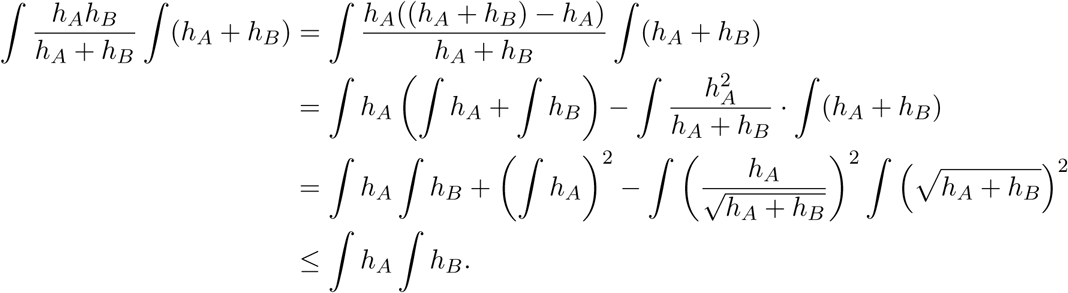

Furthermore, equality holds in (19) if and only if *f* = *γg* for some *γ ≠* 0. For the above *f* and *g*, this is equivalent to saying

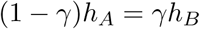

or by plugging in the definitions of *h*_*A*_, *h*_*B*_

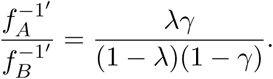

which means that *f*_*A*_ and *f*_*B*_ exhibit a constant potency ratio.

(ii) It remains to prove that (15) indeed assures the convexity of the isobole. Let any three points *P*_1_ = (*a*_1_, *b*_1_), *P*_2_ = (*a*_2_, *b*_2_) and *P*_3_ = (*a*_3_, *b*_3_) lie on the isobole. Rewrite *a*_*i*_ = *λ*_*i*_*c*_*i*_, *b*_*i*_ = (1 *-λ*_*i*_)*c*_*i*_ for 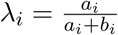 and 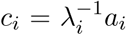, upon relabeling we may assume *λ*_1_ *< λ*_3_ *< λ*_2_. For convexity, it has to be proven that (*a*_3_, *b*_3_) lies inside the triangle Δ_1_ = Δ((0, 0)^*T*^, (*a*_1_, *b*_1_)^*T*^, (*a*_2_, *b*_2_)^*T*^). The linear transformation

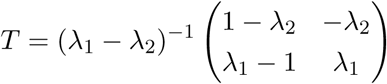

converts the ([*a*], [*b*])-plane into a ([*c*_1_], [*c*_2_])-plane, with compound drugs

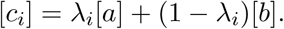

The assertion is then equivalent to 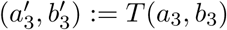 lying in Δ_2_ = Δ((0, 0)^*T*^, (*c*_1_, 0)^*T*^, (0, *c*_2_)^*T*^) = *T* Δ_1_. The dose pair 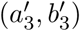 expresses the quantities of the drug compounds *C*_1_, *C*_2_ and has a ratio of

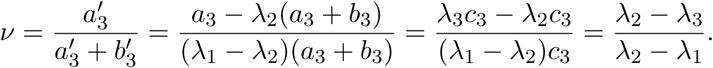

Using the associative property of the Hand model, we can apply the above derivation by replacing *f*_*A*_ = *f*_*C*1_, *f*_*B*_ = *f*_*C*2_, *f*_*AB,λ*_ = *f*_*C*1_*C*_2_,*v* and get

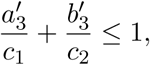

which establishes that 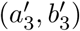 lies in Δ_2_, and thus (*a*_3_, *b*_3_) lies in Δ_1_. By this final argument we conclude convexity.

### 5.5 Derivation of the Loewe combined curve

Given *a, b* doses of *A* and *B* and the single curves *f*_*A*_, *f*_*B*_, the LOEWE predicted effect *x* has to fulfill:

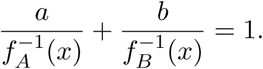

This can equally be expressed, 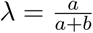, as:

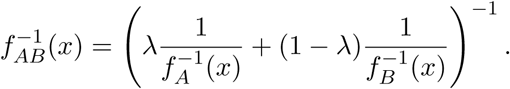

If the right hand side can be computed, then the value *x* is 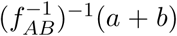. In case of Hill curves 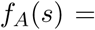 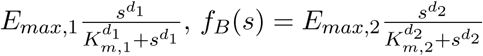, the right hand side reads as follows:

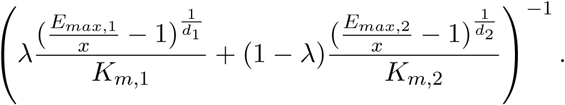

### 5.6 Derivation of the Loewe limit isobole

Let *A* and *B* be a partial and a full agent, i.e. *E*_max,*A*_ *< E*_max,*B*_. Define 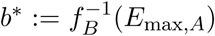. Then the Loewe model assigns an effect value to all dose pairs (*a, b*) in the domain *D* = {(*a, b*)*|b < b*^*\**^} and *E*_*L*_(*a, b*) *< E*_max,*A*_. Proof: Take an arbitrary (*a*_0_, *b*_0_) *∈ D*. Define the function *⇕*: [*b*_0_, *b*^*\**^] *→* R, *⇕*(*b*) = *f*_*A*_(*a*_*b*_) *-f*_*B*_(*b*), where *a*_*b*_ is the unique value such that (*a*_*b*_, 0), (*a*_0_, *b*_0_) and (0, *b*) lie on one line.

Then *⇕* is a continuous function and *⇕*(*b*_0_) *>* 0, *⇕*(*b*^*\**^) *<* 0. Hence for some *b ∈* (*b*_0_, *b*^*\**^) we obtain

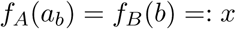

so (*a*_0_, *b*_0_) lies on the isobole at effect level *x*. Since *f*_*B*_ is increasing *x* = *f*_*B*_(*b*) *< f*_*B*_(*b*^*\**^) = *E*_max,*A*_.

On the other hand, all isoboles at effect levels 0 *< x < E*_max,*A*_ lie in the domain *D*. The isoboles at effect levels 0 *< x < E*_max,*A*_ cover the domain *D*. By continuity of *E*_*L*_ the limit isobole at the effect level *E*_max,*A*_ is forced to be the boundary of the set *D*, which is the horizontal line {(*a, b*)*|b* = *b*^*\**^}.

### 5.7 Estimate on the Error

Numerically, the ODE must not be solved starting in *x* = 0, since the simulation would result in the unfavored solution which is constant 0. If the ODE is simulated starting in *x* = *E*, the error for the dose can be approximated by

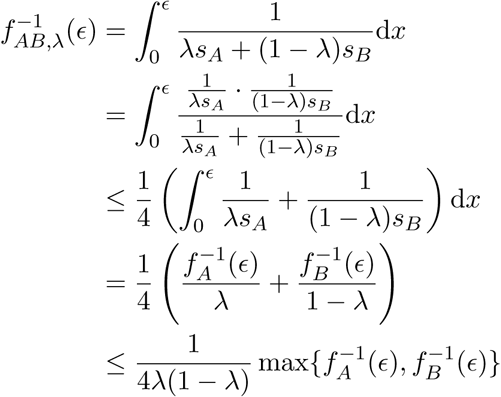

where we used the inequality of the geometric-arithmetic mean to estimate the integral. For a Hill curve *f*_*A*_ and *E* small, we have

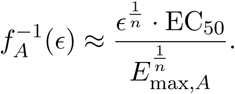

If *λ ∈* (0.03, 0.97) and ϵ= 10^*-k*^, we may thus approximate the magnitude of the error

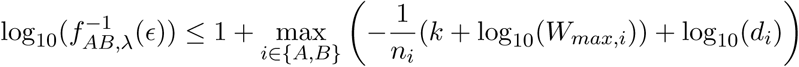

In order for the error to be less than 10^*s*^, *k* should be chosen as

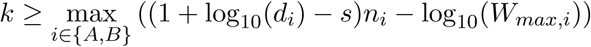

For example, in case of *n*_*A*_ = 2, *n*_*B*_ = 3, log10(*W*_*max,A*_) = log10(*W*_*max,A*_) = 5, log10(*d*_*A*_) = 3, log10(*d*_*B*_) = 2, and *s* = *-*1, one needs to choose *k* = 7.

Remark. The estimate can be improved depending on the size of *λ*. E.g. if *λ ∈* (0.3, 0.7), then

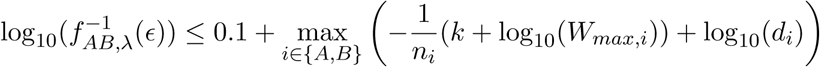

and

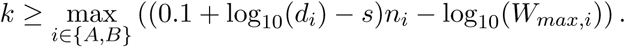

In the above example, choosing *k* = 4.3 then suffices.

### 5.8 Numerical implementation of the Hand model

The analysis of the experimental data from O’Neil *et al.* (2016) was performed in MATLAB 2017a. The code can be found on GitHub: https://github.com/ICB-DCM/NullModels

For each cell line the coefficients of the Hill curve (1) of the dose-effect curves of all drugs are fitted. We used multi-start optimization with starting values coming from a Latin hypercube and MATLAB’s lsqnonlin for the least squares fit. The BIC was used to choose between a zero response-model and the Hill curve.

The different null models were implemented in MATLAB. Equation (3), which defines the reference effect for the Loewe model, was solved by bijection. If the dose pair (*a, b*) did not lie in the set 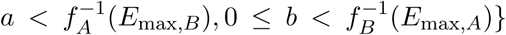, in which (3) is solvable, we extended the Loewe by 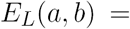 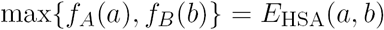. This was done since the (axis parallel) HSA isobole can be seen as continuous extension of the Loewe isobole.

The combined dose-effect curve *f*_*AB*_ of the Hand model was computed by solving the ODE (11):

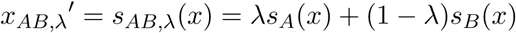

The initial condition was set to *ε* = `1e-7`, a value slightly above the ODE solvers absolute error tolerance. The MATLAB solver ode15s was used with the settings `AbsTol = 1e-8, RelTol = 1e-5` and the non-negative option.

